# A Novel Riboswitch Classification based on Imbalanced Sequences achieved by Machine Learning

**DOI:** 10.1101/2020.03.02.972778

**Authors:** Solomon Shiferaw Beyene, Tianyi Ling, Blagoj Ristevski, Ming Chen

**Author notes:** Corresponding author: Professor Ming Chen, Department of Bioinformatics, College of Life Sciences, Zhejiang University, Hangzhou 310058, P. R. China. Tel.: p86 (0)571-88206612; Fax: þ86 (0)571-88206612.

## Abstract

Riboswitch, a part of mRNA (50–250nt in length), has two main classes: aptamer and expression platform. One of the main challenges raised during the classification of riboswitch is imbalanced data. That is a circumstance in which the records of a dataset of one group are very small compared to the others. Such circumstances lead classifier to ignore minority group and emphasize on majority class, that resulting with a skewed classification. We considered sixteen riboswitch families, to be in accord with recent riboswitch classification work, that contain imbalanced dataset ranging from 4,826 instances (RF00174) to 39 (RF01051) instances. The dataset was divided into training and test set using new developed pipeline. From 5460 *k*-mers, 156 features were produced calculated based on *CfsSubsetEval* and *BestFirst.* Statistically tested result was significantly difference between balanced and imbalanced dataset (*p* < 0.05). Besides, each algorithm also showed a significant difference in sensitivity, specificity, accuracy, and macro F-score when used in both groups (*p* < 0.05). Several *k*-mers clustered from heat map were discovered to have biological functions and motifs at the different positions like interior loops, terminal loops and helices. They were validated to have a biological function and some are riboswitch motifs. The analysis has discovered the importance of solving the challenges of majority bias analysis and overfitting. Presented results were generalized evaluation of both balanced and imbalanced models, which implies their ability of classifying novel riboswitches. The scientific community can use python source code at https://github.com/Seasonsling/riboswitch, which can contribute to the process of developing software packages.

**Author Summary:** Machine learning application has been used in many ways in bioinformatics and computational biology. Its use in riboswitch classification is still limited and existing attempt showed challenges due to imbalanced dataset. Algorithms classify dataset with majority and minority group, but they tend to ignore minority group and emphasize on majority class, consequential return a skewed classification We used new pipeline including SMOTE for balancing datasets that showed better classified riboswitch as well as improved performance of algorithms selected. Statistically significant difference observed between balanced and imbalanced in sensitivity, specificity, accuracy and F-score, this proved balanced dataset better for classification of riboswitch. Biological functions and motif search of *k*-mers in riboswitch families revealed their presence in interior loops, terminal loops and helices, some of the *k*-mers were reported to be riboswitch motifs of aptamer domains and critical for metabolite binding. The pipeline can be used in machine learning and deep learning study in other domains of bioinformatics and computational biology suffering from imbalanced dataset. Finally, scientific community can use python source code, the work done and flow to develop packages.

## INTRODUCTION

Riboswitches, primarily discovered in bacteria (1), are parts of noncoding mRNA (2), predominantly present in the 5’ untranslated region (3, 4) and they have complex folded structure (5, 6). They act as a switch to transform the transcription or translation of the genes. In transcription, they turn a downstream gene ‘off’ or ‘on’ (7) in changing concentration of specific metabolites or ligands (8) and allow microbes to quickly react to change degrees of metabolites (7). A high-throughput platform showed how RNA makes structural transitions (9) kinetically compete during transcription in a new mechanism for riboswitch.

A riboswitch (50–250 nt in length) has two main classes aptamer and an expression platform (10). The aptamer region is a highly conserved domain, which is a site for binding of ligands (metabolites) and the latter one alters conformation on the binding of metabolite and hence regulates the expression of related genes (5, 6). Recently, almost over twenty diverse classes of riboswitches have been founded in bacteria, archaea (11, 12) and eukaryote but the majority in bacteria (12, 13). *Thiamine pyrophosphate* (TPP) is the only eukaryotic riboswitch, others are established in some fungi (13) for instance *Neurospora crassa*, in the case of algae and *Arabidopsis thaliana* from plants (14, 15).

In the last decades, incredible advances in big and complex omics data had emerged novel high-throughput experimental technologies such as Next Generation Sequencing (16, 17). Numerous bioinformatics databases are available to gather data for riboswitches analyses and assemble the information regarding diverse functionality of RNA molecules (18), including GeneBank (9), National Center for Biotechnology Information (NCBI), Rfam (19), Protein Data Bank (PDB), RiboD (20) and European Bioinformatics Institute (EMBL-EBI).

Many efforts have been made to develop suitable bioinformatics tools to predict the presence of riboswitches in ribonucleic acid sequences (18). Most commonly computation tools used for the analysis of riboswitches are: RiboD (20), Riboswitch finder (21), RibEx (22), RiboSW (23), mFold (24) and RegRNA (18). These available bioinformatics tools use Covariance Model (CM), Support Vector Machine (SVM) and Hidden Markov Model (HMM) algorithm. Most research exists mainly depending on the principal of multiple sequence alignment to investigate conserved sequences in already reported riboswitch. The attempt was to find out the conserved sequence of previously reported riboswitches in a targeted manner. Most reported studies are limited for the specific types of riboswitches. However, conducted frequency-dependent research revealed its importance in the classification of riboswitch (25, 26). Frequency-dependent classification uses *k*-mers counts. *K*-mers counts have many application like, building de Bruijn graphs (27) in case of *de novo* assembly from very big number of short reads, generated from next generation sequencing (NGS), used in case of multiple sequence alignment (28), and repeat detection (29).

A tremendous amount of data are generated every day that create the demand for learning algorithms that can classify, predict and analyse data more accurately (30). There are two classification categories: classification of binary format (31) and multi-class classification (32, 33).

The concept of an imbalanced dataset is defined as follows. Each family in classes of riboswitch with majority groups has more than two thousand class and minority group below thousands, which is considered as an imbalanced sequence. Whereas, the imbalanced group used and treated with Synthetic Minority Over-Sampling Technique (SMOTE) and thereafter it is called a balanced dataset. The classification with imbalanced data gives favors for a sample with the majority class (30). Imbalanced data occur as a circumstance where the records of a dataset of one class are very little regarding the other classes’ dataset. This leads classifier algorithms to ignore minority groups and emphasize on majority class, which can result in skewed accuracy of the classifier. The value of the accuracy of the classifier might be high, but minority class misclassified. Several findings have been done for riboswitch classification (25, 26) based on imbalanced data. However, data resampling can be a solution to handle the class imbalance problems (30). Synthetic Minority Over-Sampling Technique (SMOTE) has been discovered in 2002, which is a sampling-based algorithm. Synthetic Minority Over-Sampling Technique (34) balances the class distribution of an imbalanced dataset through an incrementing approach on some virtual samples.

To address the needs for riboswitch prediction, conserved nucleotide frequency counts are considered. SMOTE was used for resampling. Different machine learning algorithms are used for evaluation such as: Random Forest (RF), Gradient Boosting (GB), Support Vector Machine (SVM), K-Nearest Neighbors (KNN), Na*ï*ve Bayes (NB) and Multilayer Perceptron (MLP). The performances of each algorithm on classification were derived from the confusion matrix, which reveals the number of matches correctly and mismatched instances of riboswitches. Specificity, sensitivity, accuracy, and macro F-score were calculated. That parameters are the main performance evaluation criteria for machine learning algorithms (35–38).

## RESULTS

### Features engineering and selection

Riboswitch families considered for this analysis and their corresponding details were presented and analyzed in Figure 1 and features where clustered (Figure 2). Looking into instances in riboswitch, there were differences in representation between families range in distribution from Cobalamin riboswitch (4,826) to PreQ1-II (39). Out of 16 riboswitch class, Cobalamin riboswitch, TPP riboswitch (THI element), and Glycine riboswitch contributed for 68% and the remaining 13 riboswitch family has 32% instances. The performances of algorithms and methods were computed and evaluated based on training and test set (details in the methodological approach part). We produced 5460 *k*-mers and then calculated 156 features which include *k-*mers up to six, which was consistent with previous research (26) based on *CfsSubsetEval* and *BestFirst* (Figure 3).

**Figure 1.**
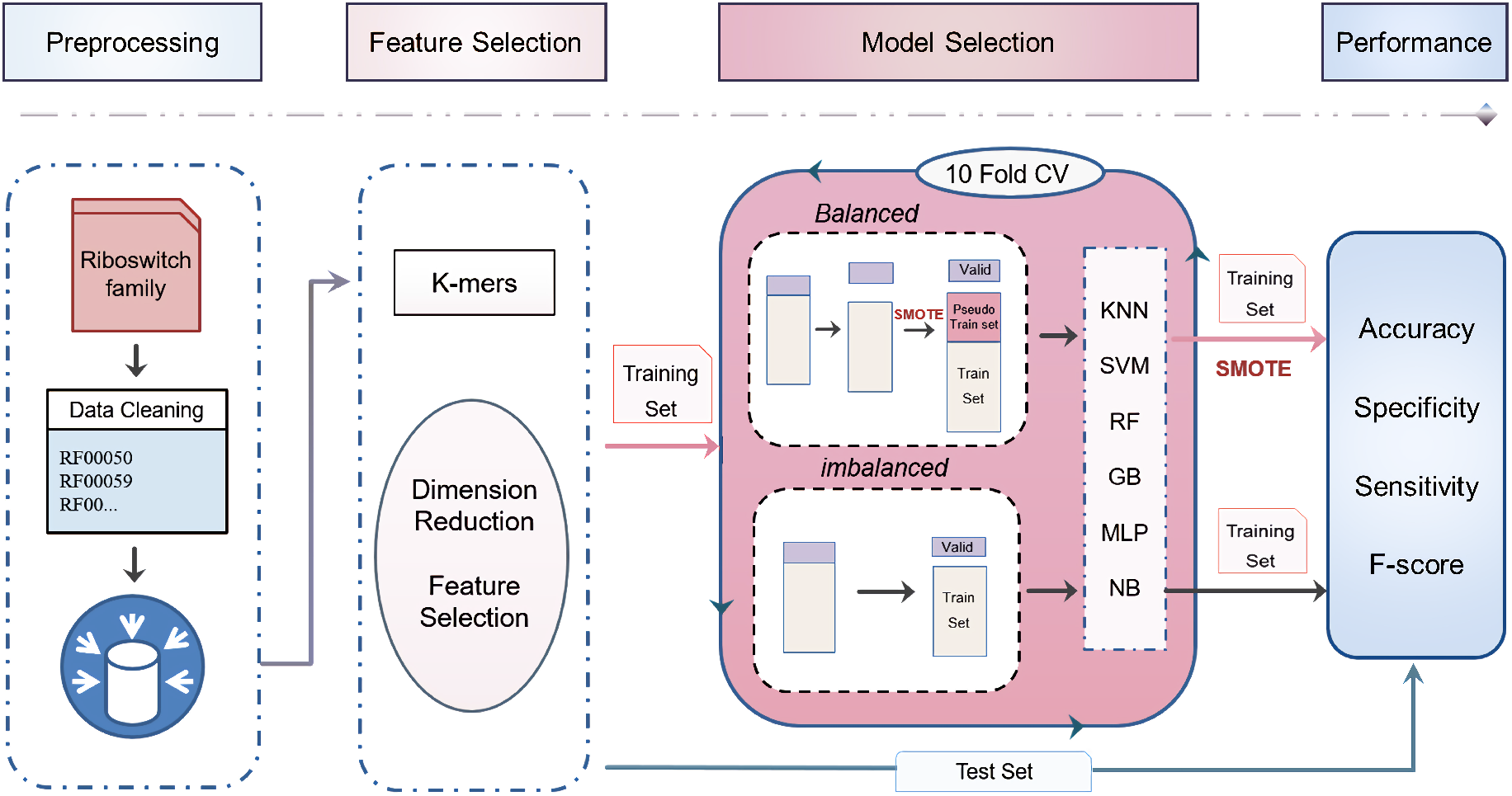
The workflow used to analyze imbalanced and balanced datasets to compare the computational performance of machine learning algorithms for classification.

**Figure 2:**
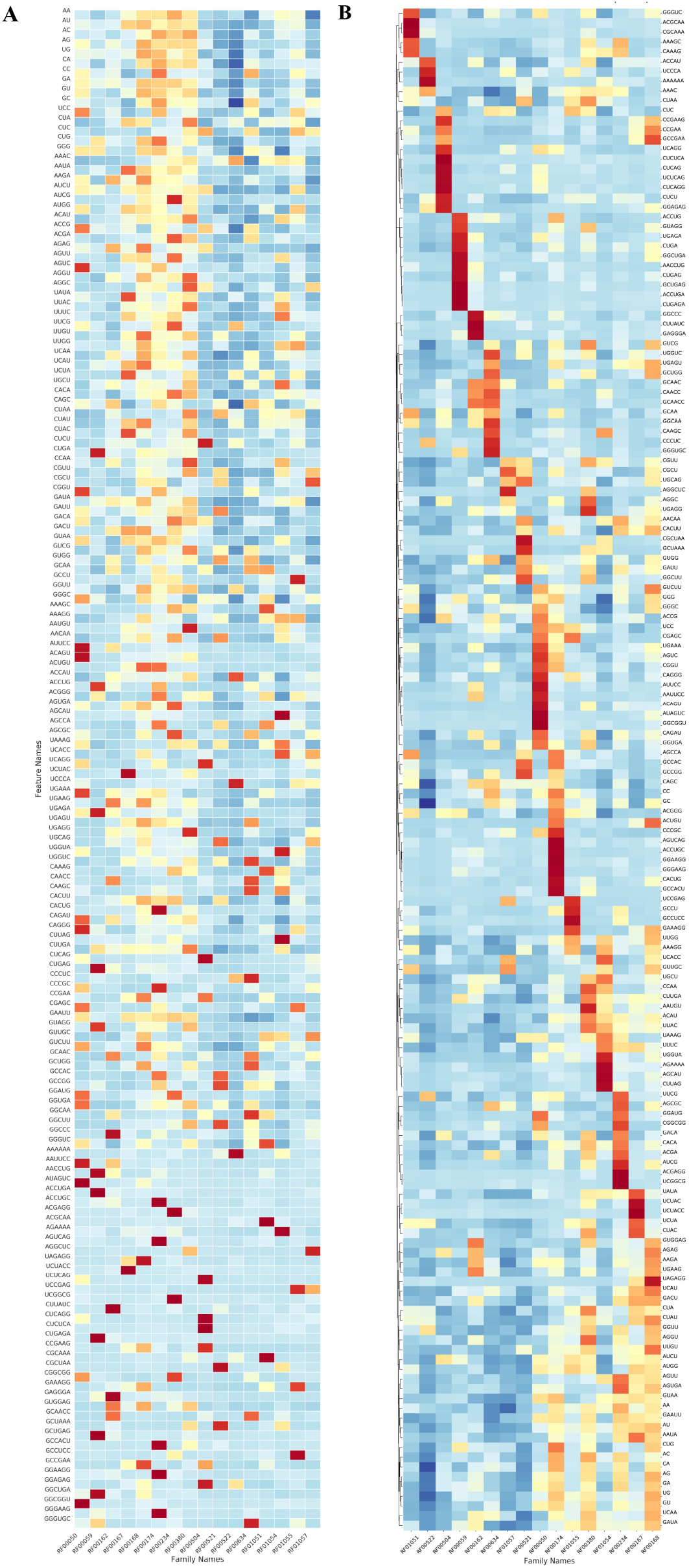
Heat-map in this figure represented as A) row-normalized *k*-mer counting distribution, rows correspond to the *k*-mers, and columns revealed 16 classes of riboswitch and B) the clustering heatmap depicts feature clustering, clustered features were important for classification in that family. Red means a high relatively counting number while blue means lower.

**Figure 3.**
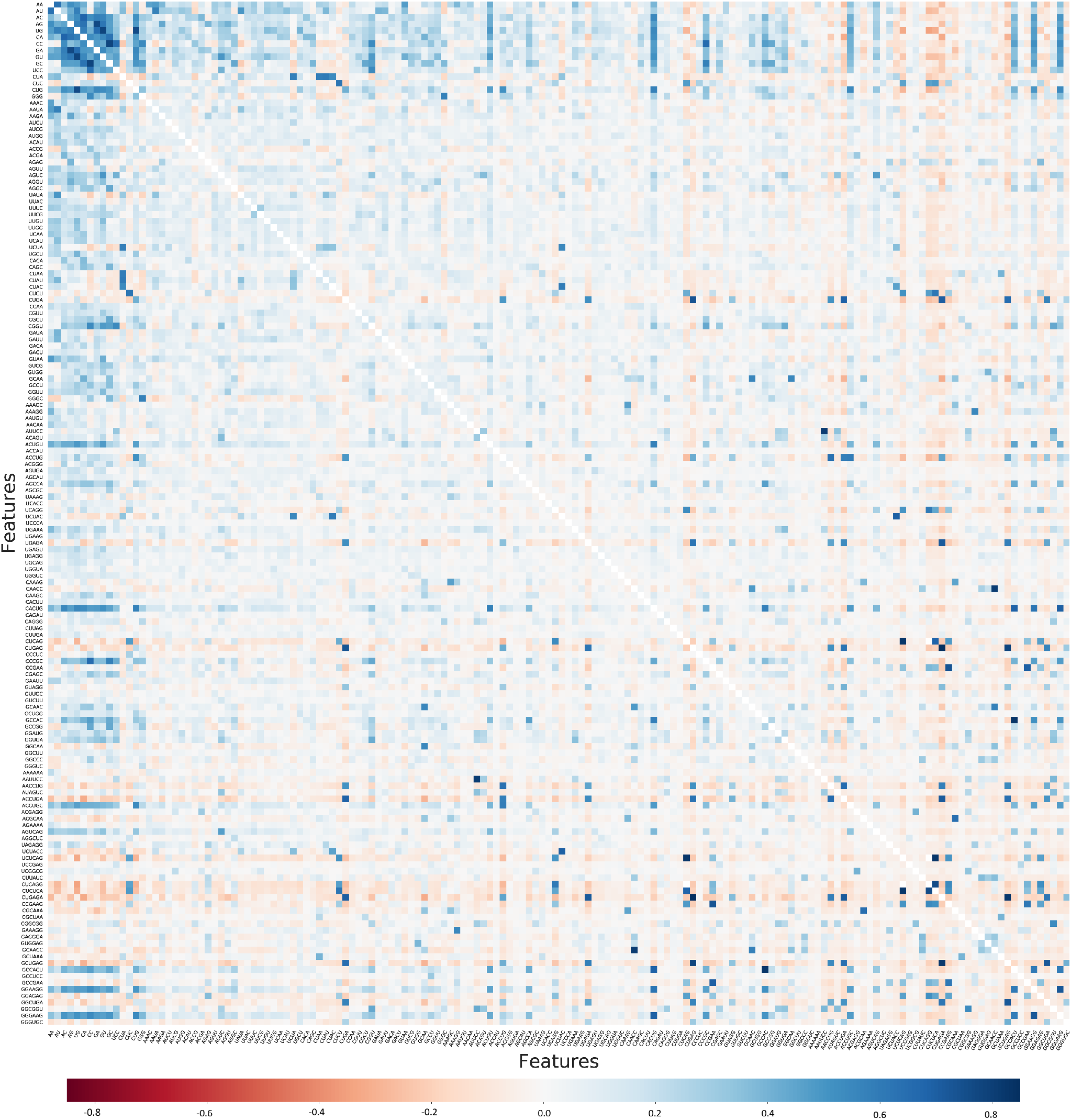
Heat-map showed 156 feature correlation, the diagonal white line represented their correlation value equals to one. Blue means a positive correlation while red means a negative correlation.

### Imbalanced class on classification performance

Classifiers on minority class resulted in F-score value from 0.50 (NB) to 0.94 (MLP), while on majority class the range is from 0.91 to 1.00, as indicated in Table 1 and Figure 4. Riboswitch families considered for classification was present in Supplementary Table S1. The average performance of each classifier is computed using mean and standard deviation for parameters: accuracy, specificity, sensitivity and F-score.

**Table 1.**
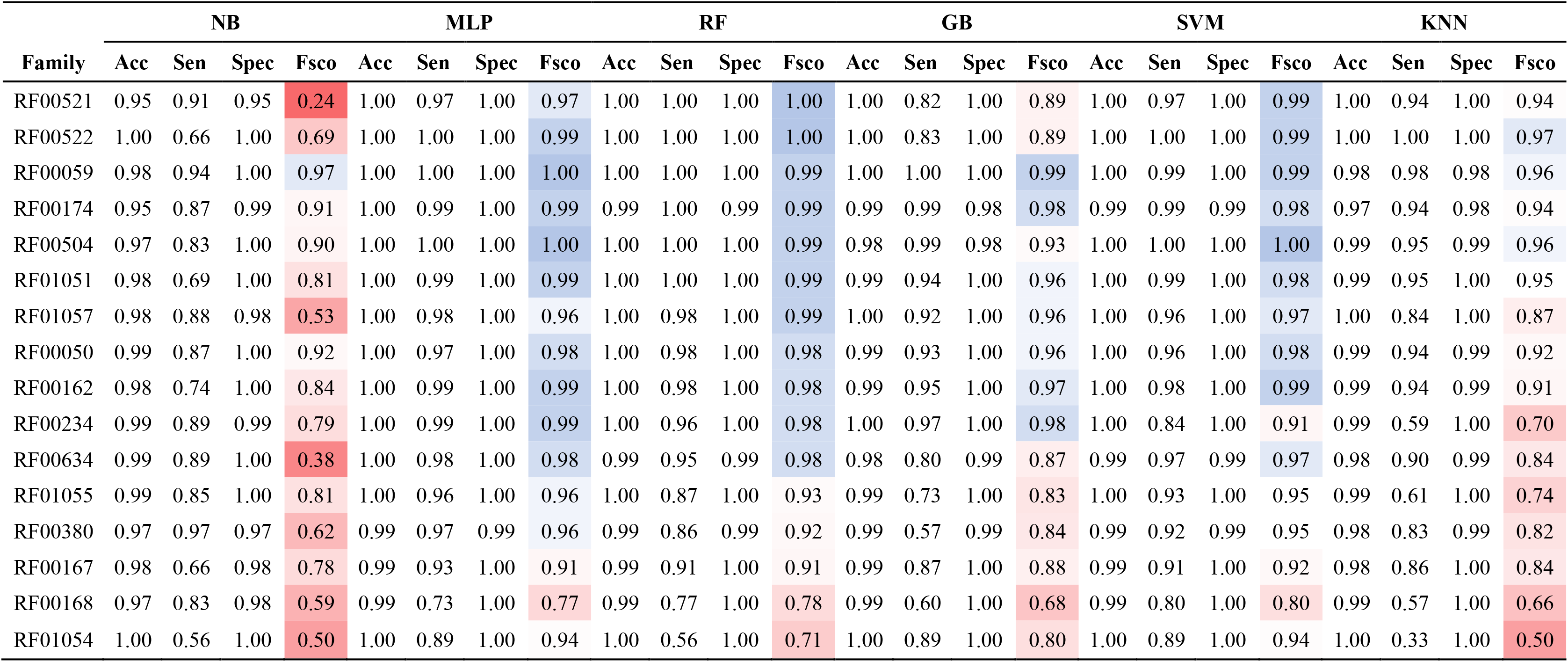
Accuracy, sensitivity, specificity and F-score parameters used for Naïve Bayes, Multilayer Perceptron, Random Forest, Gradient Boosting, Support Vector Machine and K-Nearest Neighbors algorithms evaluation when applied on the imbalanced dataset. The color trend of F-score from blue to red indicates performance from the best to the poorest.

**Figure 4.**
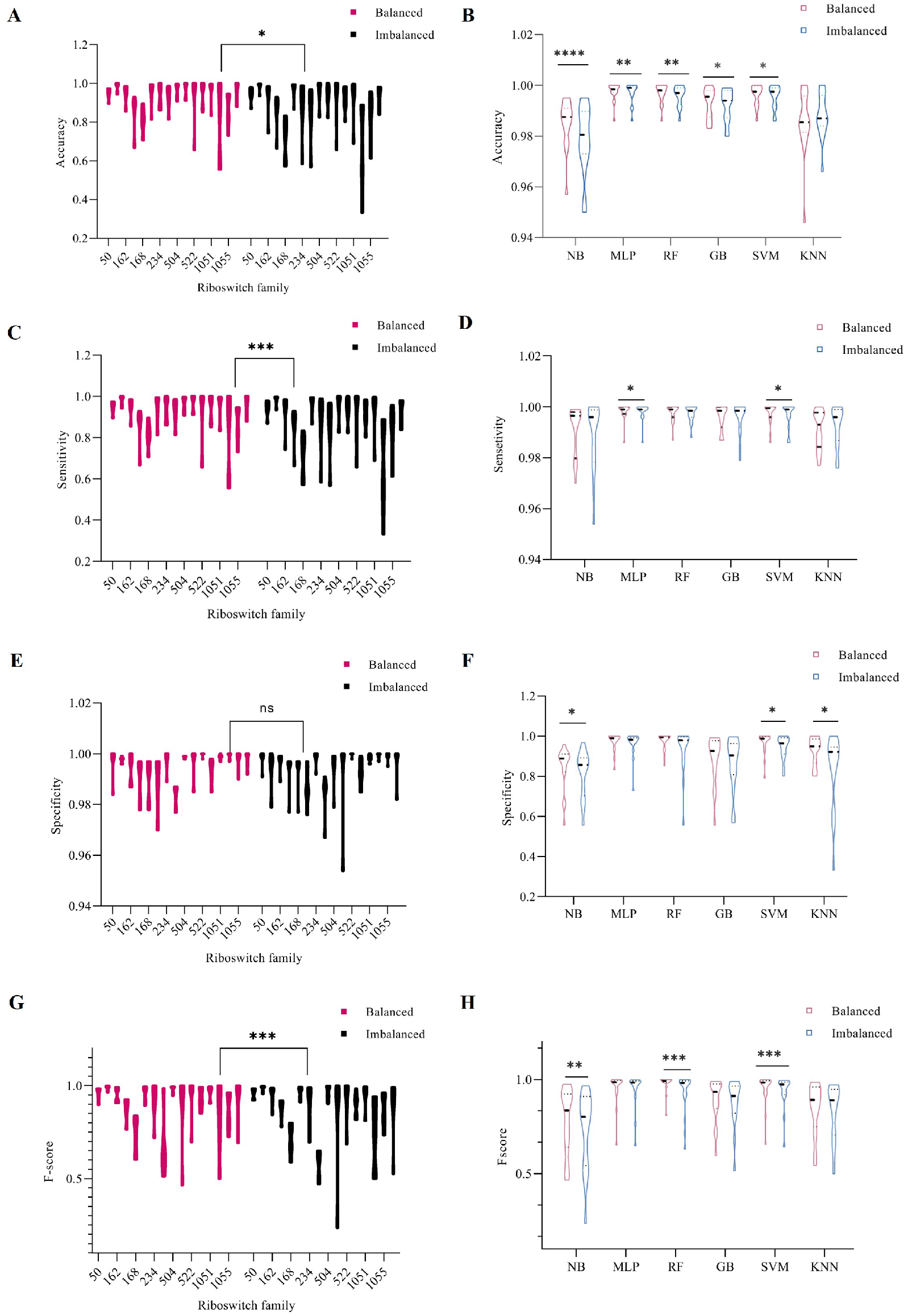
The figures showed a comparison of the balanced and the imbalanced datasets using the Wilcoxon rank test, ACEG) showed performance evaluation of classifiers on both dataset and BDFG) Performance of selected six machine learning algorithms. Violin box was used to depict the statistical differences between two group were provided as the plots. (* indicated significant difference of p < 0.05, ** denoted very significant difference of p < 0.01, and *** showed very significant difference p < 0.001.

The comparative analysis of six algorithms has revealed that MLP performs best, while NB performed the poorest results (Supplementary Table S2). RF00174, RF00059, RF00504, RF00522 classified better than others with minority classes like RF01054, RF00634, RF00380 (Table 1). F-scores of MLP and RF for the majority group (RF00174) were 0.997 and 0.996, respectively. In the minority group, classifiers with high accuracy had F-score up to 0.50 in the case of NB. The computed minimum value in overall NB analysis in RF01054, RF00634, and RF00521 were 0.50, 0.38, and 0.24, respectively. Accuracy of all algorithms across all riboswitch family showed value greater than 0.97. In confusion matrix, predicted family and true family exhibited performance of classifiers and riboswitch classification (Figure 5).

**Figure 5.**
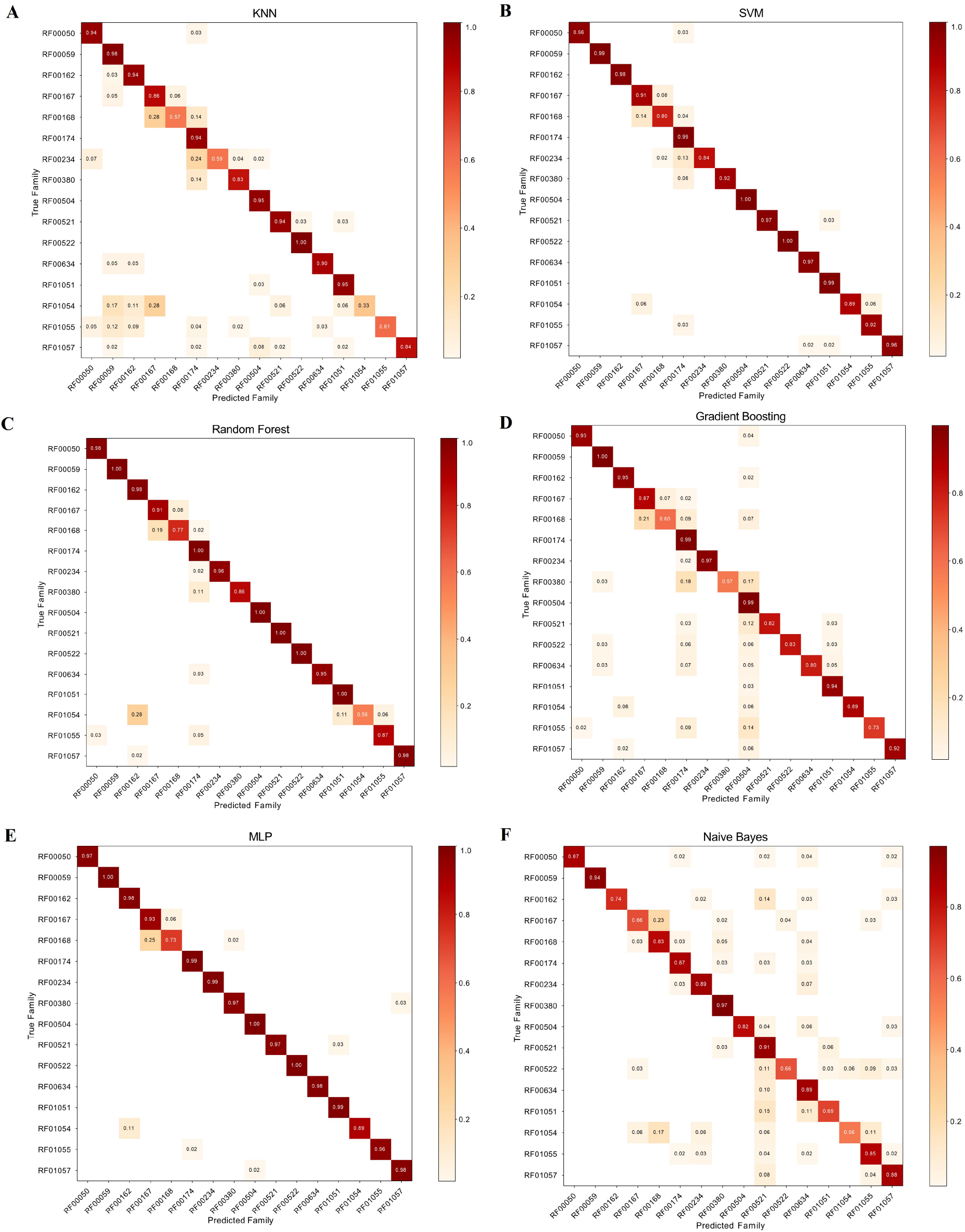
Confusion matrix for the imbalanced dataset from independent test experiments depicted True family and Predicted family with classifiers as: A) K-Nearest Neighbors, B) Support Vector Machine, C) Random Forest, D) Gradient Boosting, E) Multilayer Perceptron and F) Naïve Bayes.

### SMOTE balancing on classifiers performance

The overall analysis computed for frequency counts of all family had discovered improved performances of classifiers (Supplementary Table S2, Table 2 and Figure 4). RF00059 and RF00174 results showed F-score between 0.93 and 1.00. In the case of NB and KNN the results of the F-score indicated their poorer performance with a value less than 0.84. Performance evaluations have revealed that KNN, NB, SVM, MLP, RF and GB can be used for classification of riboswitch (Figure 6).

**Table 2.**
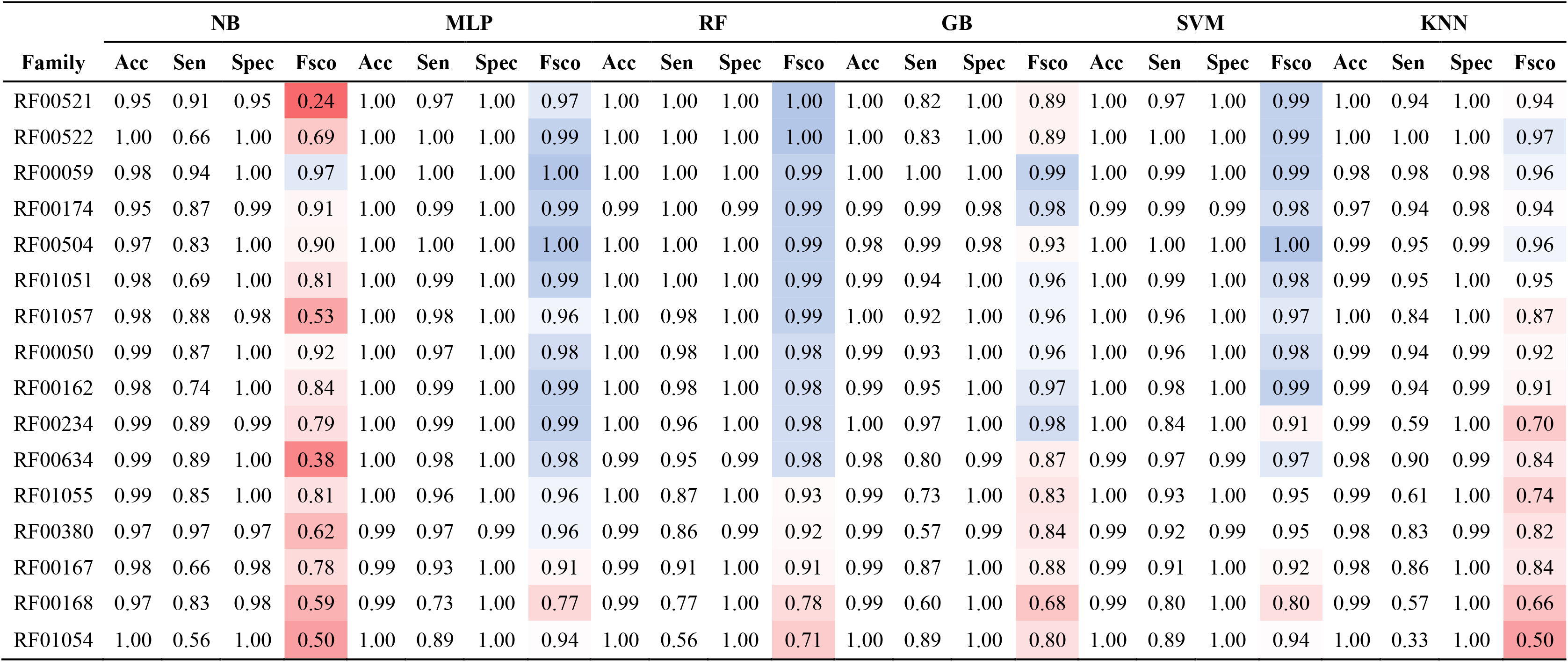
Performances of Naïve Bayes, Multilayer Perceptron, Random Forest, Gradient Boosting, Support Vector Machine and K- Nearest Neighbors were evaluated using the balanced dataset from 16 riboswitch families measured by using accuracy, sensitivity, specificity and F-score. The color trend of F-score from blue to red indicates performance from the best to the poorest.

**Figure 6.**
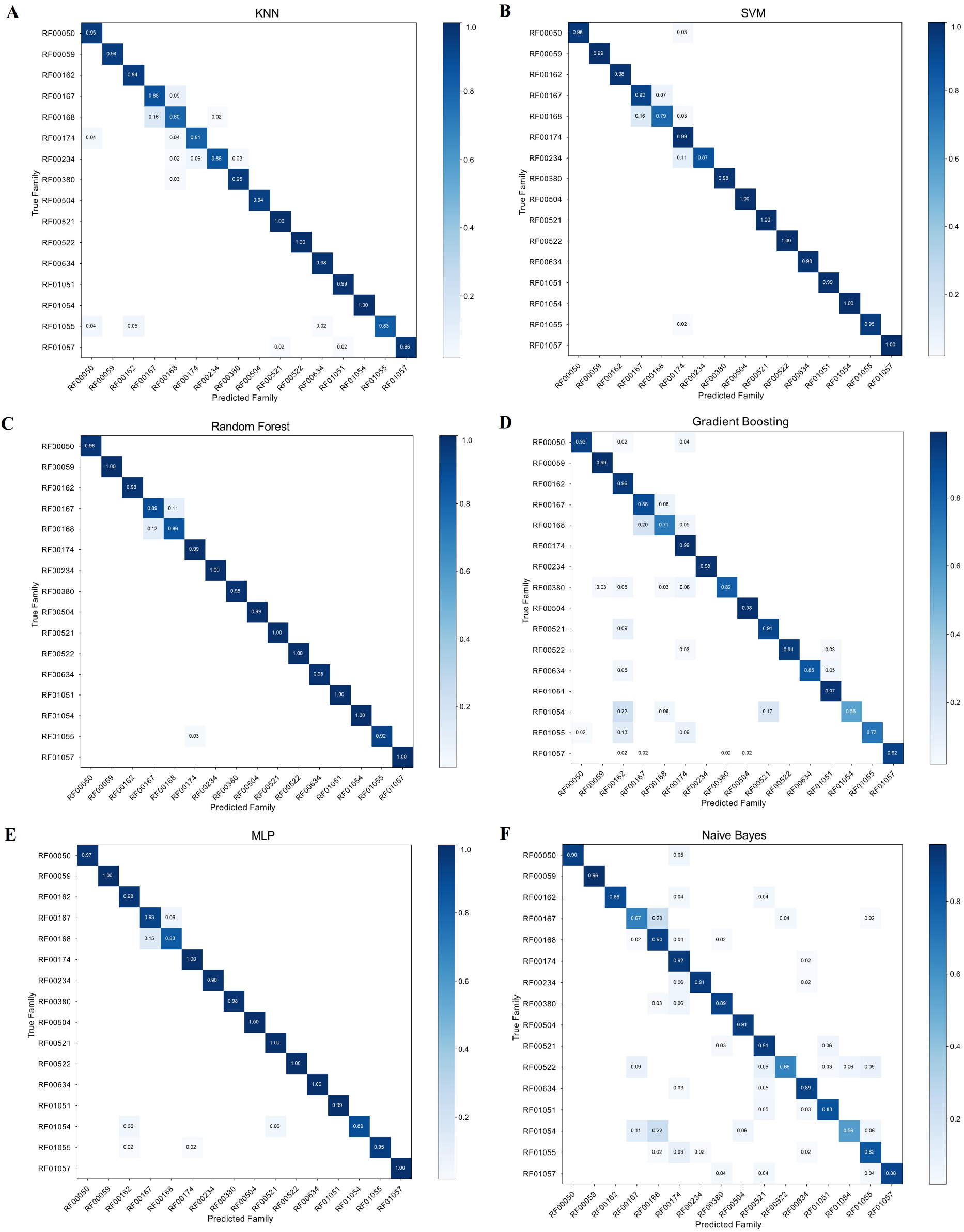
Confusion matrix for the balanced dataset from independent test experiments showed True family and Predicted value with classifiers as: A) K-Nearest Neighbors, B) Support Vector Machine, C) Random Forest, D) Gradient Boosting, E) Multilayer Perceptron and F) Naïve Bayes.

As presented, Random Forest and MLP exhibited the consistently higher accuracy and F-score values compared to NB, GB, SVM and KNN. Figure 4 and Table 2 have shown that SMOTE improves riboswitch classification and algorithm performances.

The overall accuracy of SMOTE analyzed dataset (balanced dataset) showed consistent and better results than an imbalanced dataset (Table 1, 2 and Supplementary Table S2). The specificity of NB, MLP, RF, GB, SVM and KNN were better in the balanced dataset than when applied on an imbalanced dataset. Calculated sensitivity results were slightly better in balanced instances. Surprisingly, evidence discovered in that F-score value in all the models showed that a balanced dataset can improve the classification of riboswitches. Balanced dataset not only increased classification accuracy but also algorithms performances. Table 2 has depicted F-score values increasing from 0.50 while in the case of the imbalanced dataset to 0.84.

### Application of statistical significances

Statistical computation using the Wilcoxon rank test (39) between balanced and imbalanced datasets depicts significant differences between these two groups. In addition, the performance of NB, MLP, RF, GB, SVM and KNN statistically showed variation in accuracy, specificity, sensitivity and F-score values. Statistically very significant differences were noticed between balanced and imbalanced in F-score and specificity (*p* < 0.001) and accuracies were significantly different (*p* <0.05), whereas sensitivity showed no significant difference between the two groups (Figure 4, Supplementary Table S3).

SVM was a very significant difference in all parameters used for performance evaluation, F-score (*p*<0.001) whereas accuracy, specificity and sensitivity were significantly different (*p*<0.05). RF performance in both groups has shown very significant differences in F-score (*p*<0.001) and accuracy (*p*<0.01) (Figure 4 and Supplementary Table S2). In KNN we did not notice statistical significant differences in all used parameters, except significant differences in specificity (*p* < 0.05).

MLP of the balanced and imbalanced group depicted very significant differences in accuracy and sensitivity (p < 0.01). GB showed significant differences only in accuracy (p<0.05). Finally, both imbalanced and balanced datasets in the case of NB have shown very significantly differences in F-score (p<0.01), accuracy (p < 0.001), whereas specificity was a significant difference (p<0.05). Accuracy of all classifiers is significantly different at different levels in both groups except in KNN (Figure 4 and Supplementary Table S3).

### Biological functions of clustered *k-mers*

*K-mer*s counting was extracted from distribution heat-map (Figure 2A and 2B), which depicted feature clustering and high relative count number. These clustered *k-mer*s were used for biological function and motif searching. Accordingly, in Table 3 riboswitch families and their *k-mer*s were used to verify their biological functions. Structural analysis from *k-mer*s coverage results is depicted in the case of RF00174 (A) and RF01055 (B). In every individual base, the color gradient scale indicates a normalized count. Results depict different color scale in each region and their interior loops, helices, and terminal loops (Figure 7).

**Table 3.**
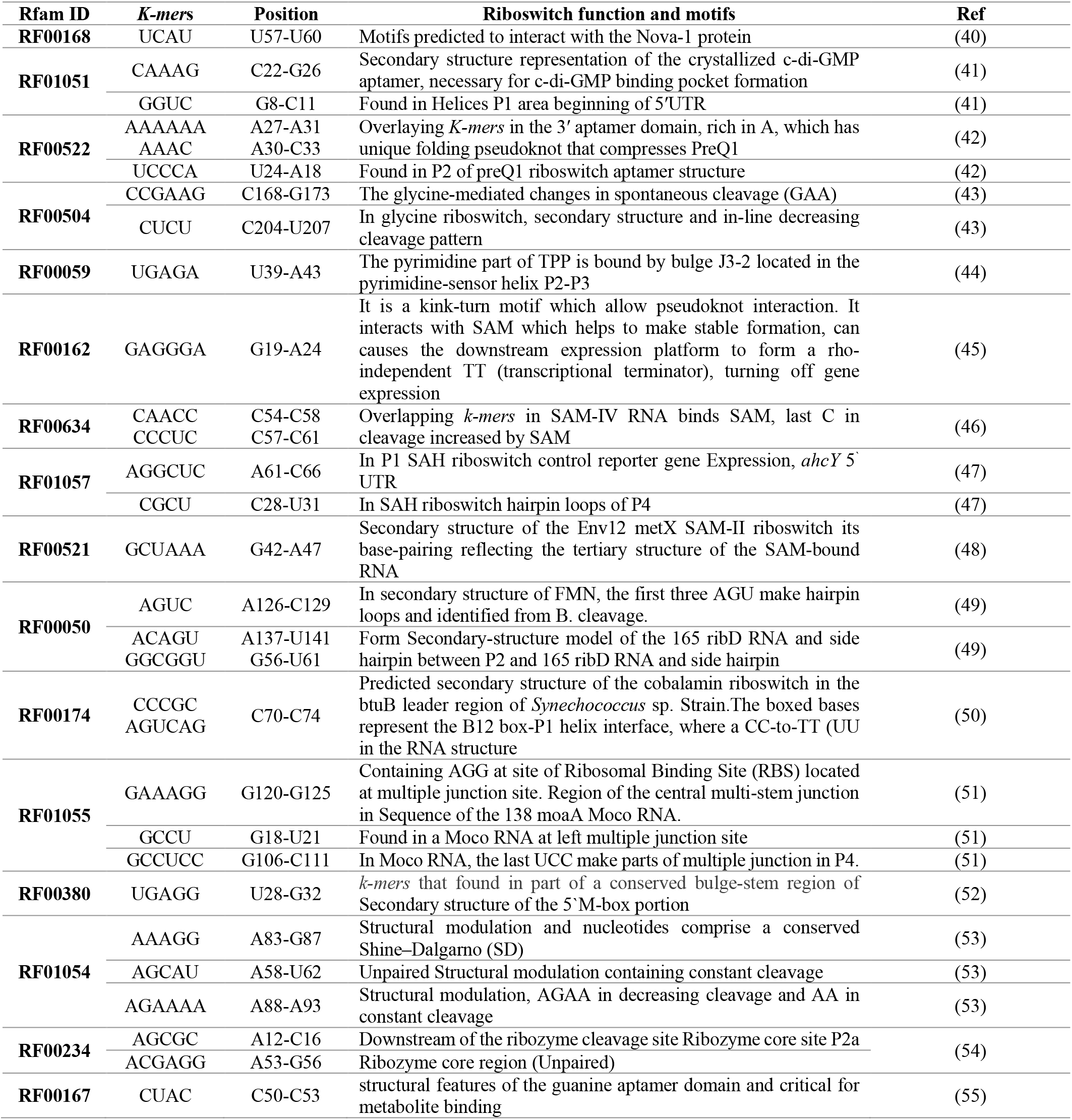
Clustered *k-mers* from Figure 2A and 2B used for validation of their biological function and reported riboswitch motifs. Nucleotide location designated refers to match with their position reported in reference.

**Figure 7.**
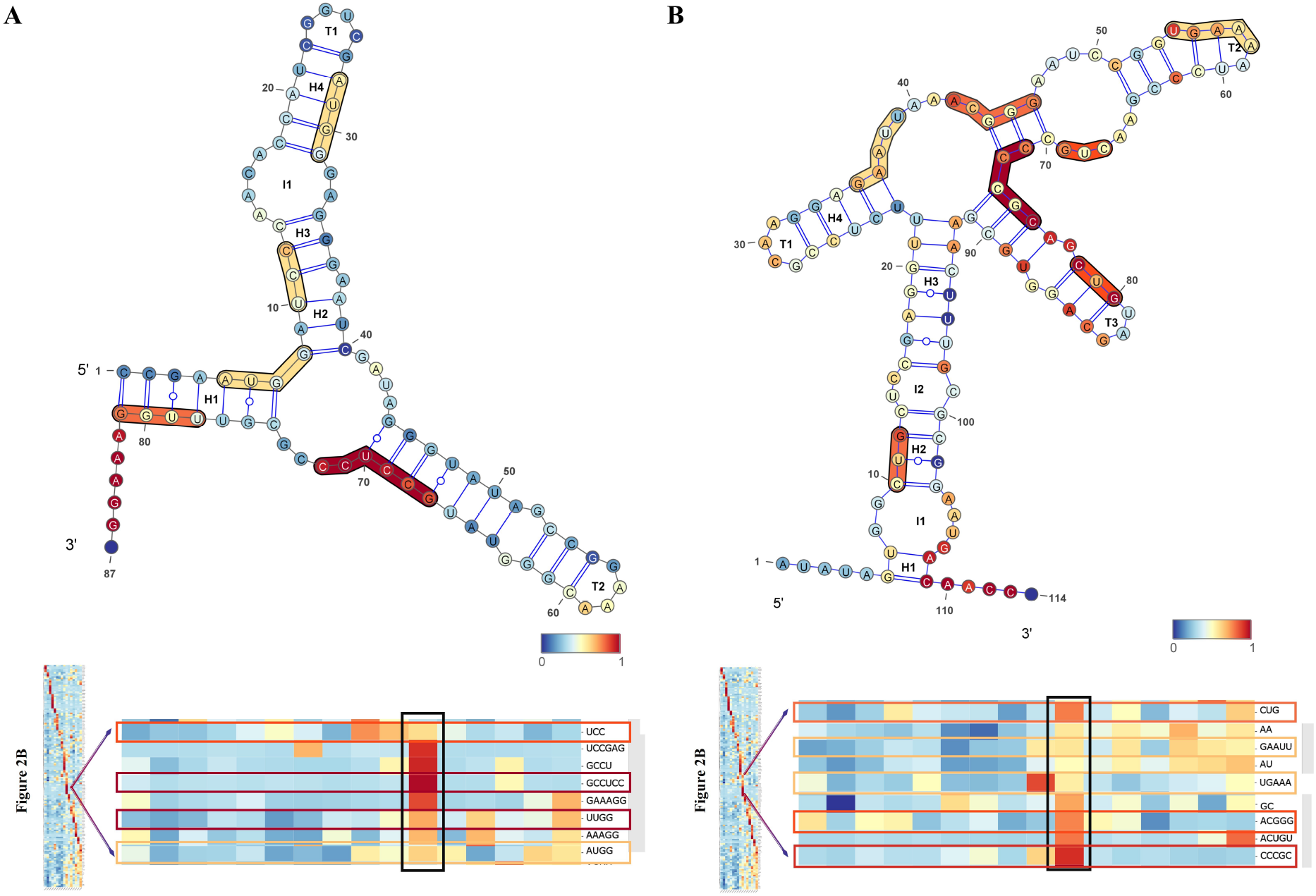
Secondary structure of RF00174 Cobalamin riboswitch (Acido bacterium sp. MP5ACTX8) (A) and RF01055 MOCO riboswitch class (B). In every individual base, the color gradient scale represents a normalized hit number from 156 features aligned to the sequence. The different color scale in each region represent its coverage of the k-mers in the family that it represents. Whereas, I, H, and T are abbreviations for Interior loops, Helices, and Terminal loops, respectively.

## DISCUSSIONS

Machine learning has an enormous capacity to boost our knowledge in the classification of riboswitch, an area that is still in the early stage of a comprehensive investigation. Numerous machine learning applications have been developed based on different methods to detect riboswitch. However, most riboswitch classification studies applied machine learning algorithms on the imbalanced dataset (25, 26). Several findings revealed the impact of an imbalance dataset on correct classification and performance of algorithms (25, 26, 30). Chawla and colleagues proposed SMOTE method of treating imbalanced datasets for better classification of majority and minority instances (30, 34). SMOTE based balancing of dataset improves the oversampling minority classes accurately and also produce a dataset that does not influence majority class.

In this analysis, there are imbalances of instances in the riboswitch family. Comparative results revealed the reality of the impact of such imbalance on classification which has widely been reported (Supplementary Table S1 and Table 1). Imbalanced distribution exhibited variation from 4826 majority class (Cobalamin riboswitch) to 39 minority class (PreQ1-II riboswitch). In this type of circumstances general classifiers, when encountering such imbalanced data, favor class with majority instances (30, 34). The analysis also revealed in imbalanced and balanced confusion matrix the same problem (Figure 5 and 6). Out of 16 riboswitch class, cobalamin riboswitch, TPP riboswitch (THI element), and glycine riboswitch sum up contribution was 68% and the remaining 13 riboswitch family has 32% instances. In Table 2, full sequences grouped into two sets training (70%) and test set (30%) was selected and performances of classifiers were evaluated regarding sensitivity, accuracy, specificity and F-score. The correlation heat-map in Figure 3 indicates the relationships between *k*-mers.

Dataset of imbalanced sequences in riboswitch showed different performances of classifiers ranked as: MLP - the best and NB – the poorest regarding their mean scores that range from 0.771 to 0.961. In Table 2, individual score results of this method have shown best result in RF00234, RF00522, RF01057 (1.00 in RF): greater values than reported in other study using BLAST^+^ (26, 56), which is most popular tools in analysis of sequence similarity (56) and others (25, 26). Conversion of sequences into vector revealed good results in both groups used for analysis (Table 1, 2 and Supplementary Table S2). In protein study, protein sequence converted into feature vectors showed good performance in cases of SVM and KNN (57–60). RF00174, RF00059, RF00504, RF00522 predicted better than others with minority classes like RF01054, RF00634, RF00380 (Table 1 and 2). The class with maximum instances (RF00174) resulted in an F-score value greater than 0.94 in all classifiers except NB, which had a value less than 0.93 in both cases.

NB classifier depicted poor performance in imbalanced dataset compared to other classifiers and its accuracy, sensitivity, specificity, and F-score had the following values 0.979, 0.989, 0.814 and 0.705, respectively (Table 3). These results were improved to 0.985, 0.991, 0.841 and 0.771 when datasets were balanced. Compared with the F-score value reported by Hugo and colleagues (NB-HEXCFS-0.525), the changes indicated the influence of imbalanced datasets on the performance of classifiers. Similarly, improved performance of NB on large datasets has been reported (61).

Table 2, Supplementary Table S2, and Figure 4 indicate that the proposed method of balancing instances increases classifier performances. The used approach was also reported as a solution for machine learning (62). RF shows the best result followed by MLP, which revealed optimal performance. On the other hand, Naïve Bayes has poor performances in imbalanced dataset classification, which is in accordance with Mwagha and colleagues (63, 64). The overall comparison revealed that balanced datasets are better for classification of riboswitch, their performances were compared to BLAST^+^ (26) and other finding (Table 1, 2 and Supplementary Table S3).

The *k*-mers position in the secondary structure illustrated riboswitch biological function and motif (Table 6 and Figure 7). In RF00174, *CCCGC k*-mers had predicted the secondary structure of the cobalamin riboswitch in the btuB leader region of Synechococcus. In cases like RF00168, *AAAAAA k*-mer had motifs predicted to interact with the Nova-1 protein, overlaying *K*-mers in the 3′ aptamer domain, rich in A, which has unique folding pseudoknot that compresses PreQ1 (Scott et al. 2016). Turning off gene expressing observed in RF00162 with *GAGGGA k*-mer, is a kink-turn motif which allows pseudoknot interaction. It interacts with SAM which helps to make stable formation, and can cause the downstream expression platform to form a rho-independent TT (transcriptional terminator), turning off gene expression (Montange and Batey 2006). Overall, *k*-mers and their biological function for this study are summarized and described in Table 3.

The pipeline can be used in machine learning and deep learning study in other domains of bioinformatics and computational biology that suffer from an imbalanced dataset. Finally, the scientific community can use the python source code for analysis of interest as well as to develop suitable software packages.

## METHODOLOGY

We showed a complete evaluation of different machine learning approaches for classification and predicting regulatory riboswitches. First of all, we present the benchmark datasets and data mining approach followed by feature engineering that were done through testing. In addition, model selection methods were used to model and compare balanced and imbalanced dataset problem (65). These methods are implemented in an open-source machine learning platform called WEKA 3.8.3 (66, 67), SMOTE [31] and Python 3 (68), which allow evaluating different parameters and algorithms for classification and prediction of the riboswitch. Lastly, we described the results of classification from the learned models. The workflow for the analysis of imbalanced and balanced datasets used for performance evaluation of different machine learning algorithms found in Figure 1. This workflow can be used for other research area that suffers from challenges of imbalanced dataset. All datasets used Python source codes, which are available at https://github.com/Seasonsling/riboswitch.

### Data preprocessing

The dataset for the investigation was collected from Rfam 13.0 (19) and other datasets that were already produced (26) for comparison of our new methods. Rfam is a source that collects RNA families including riboswitch (19). There is a need to use a machine learning approach to train algorithms to classify riboswitch as it has been happening in other areas of bioinformatics. Only 16 families have been used to compare with previous research work and they clearly show the impact of an imbalanced dataset on the performance of classifiers. Preprocessing, cleaning and filtering were done, as well as handling missing values, noisy data, redundant features and irrelevant features to affect the accuracy of the model (67). The datasets that contain sequences per family are shown in Supplementary Table S1.

### Feature selection

FASTA format dataset was used for *k*-mer (*1*≤*k*≤*n*) frequency counts through executing in the R package called *kcount* (69). In order to obtain a sufficiently informative *k*-mer counting matrix for the task (70), we set *k* value to 6, and finally got 5,460 features, which was consistent with some other researchers (26). This *k-mers* composition was used to make frequencies of each riboswitch. This avoids unnecessary computing power consumption and dimensional disaster caused by extremely sparse matrices due to high *k* values as well.

Attribute evaluators *CfsSubsetEval* and *BestFirst* were used for dimensionality reduction and searching of the space of attribute subsets by greedy hill-climbing augmented with a backtracking facility (71). WEKA 3.8.3 was used to implement the task (66, 67). Feature selection was done for the dimensionality reduction and thus for decrease processing load (72, 73).

### Imbalanced data

The dataset for this finding contains the imbalanced datasets ranging from 4,826 instances (RF00174) to 39 (RF01051) instances (Supplementary Table S1). Learning from the imbalanced datasets that become critical concerns nowadays, particularly when minority class contains small instances in its dataset (25, 26, 74). Mainstream methods dealing with imbalance data can be roughly divided into two categories. The first category considers the difference in the cost of different misclassifications (75), while the second one mainly focuses on training data sampling strategies. Here over-sampling and under-sampling were common techniques used to adjust class distribution. However, traditional random oversampling adopts the strategy of simply copying samples to increase the minority samples, which is prone to the problem of overfitting that makes the information learned by the model too special and not generalized (76).

SMOTE improved scheme based on random oversampling was applied (59). The basic idea of the SMOTE algorithm is to analyze a small number of samples and to add new samples to the data set based on a small number of samples.

The used algorithm flow is as follows:

For each sample *x* in a few classes, calculate the distance from all samples in a few samples sets by Euclidean distance, and get its *k*-nearest neighbors.

Set a sampling ratio according to the sample imbalance ratio to determine the sampling magnification *N*. For each minority sample *x*, randomly select several samples from its *k*-nearest neighbors, assuming that the selected neighbor is 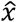.

For each randomly selected neighbor 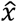, construct a new sample with the original sample according to the following formula:

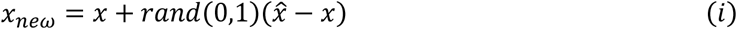

SMOTE was deployed through importing “imblearn.over_sampling” module in Python 3 and it was applied both in 10-fold cross-validation and building final model processes, as shown in Figure 1.

### Machine learning models

A crucial step in machine learning is model selection, as the performance of algorithms is sensitive to the calibration parameters. Configuration and choice of the hyper-parameters are found to be crucial. For our dataset, we calibrated a model using 10-fold cross-validation. Firstly, the complete feature selection of *k-mers* dataset was divided into two parts randomly: 70% of data were training set, while 30% of data were test set. The 70% training set was used to build multiclass classification models and determine the hyper-parameters through 10-fold cross-validation. Then, the test set was used to test the final generalization performance of the balanced and imbalanced models. In order to increase the credibility of comparison results and to ensure the repeatability of the results, all datasets were chosen randomly. Input data and model parameters except for the step of SMOTE processing were strictly consistent for both balanced and imbalanced models. This task was left to make *pipeline* module and *Pipeline* object in Python package *imblearn* (0.5.0), which ensures that in cross-validation or generalization testing, SMOTE only treats the training data used to build the cross-validation model or the final model. By this means, the validation set in each fold cross-validation was consistent in all models just as in the case of 30% test set.

### Experimentation classifiers

Random Forest is commonly used machine learning algorithm (77) with different successful function in computing and bioinformatics (77–79). It randomizes the variables (columns) and data (rows), generating thousands of classification trees, and then summarizing the results of the classification tree. In this research, the mean decrease impurity method was used.

SVM is a simple and efficient method for solving the quadratic programming problem (80) through computing the maximum marginal hyper-plane (66). In SVM, the kernel function implicitly defines the feature space for linear partitioning, which means the choice of kernel function is the largest variable of SVM.

Gradient boosting is a boosting algorithm, which belongs to ensemble learning as well as random forest and proved to have great performance in imbalance problem. It builds the model in a stage-wise fashion, and generalizes them by allowing optimization of an arbitrary differentiable loss function (81).

Another classifier is *k*-Nearest Neighbors (KNN) which also named IBK (instant-based learning with parameter *k*). This classifier offers numerous choices to speed up the undertaking to locate nearest neighbors (67), NB (Naïve Bayes) classifier based on Bayes’ theorem [49]. This is a probability-based model in the Bayesian networks (82). MLP are commonly used machine learning algorithms (83). ncRNA classification and prediction problems have been widely conducted based on the six selected algorithms for this analysis (84–86) and riboswitch classification and prediction (3, 26).

The tuning of KNN, SVM, RF, GB and MLP was carried out on the training set by evaluating the macro F-score in Python 3. The configurations of their parameters are as follows:

KNN: number of *k* = *{2, 4, 6, 8, 10, 12, 14, 16}*
SVM: type of kernel function = *{linear, poly, rbf, sigmoid}*
RF: with the method of GridSearchCV and kfold = *10*, the number of trees in the forest = *{500, 1000, 2000}*, the maximum depth of the tree = *{10, 15, 20}*
GB: with the method of GridSearchCV and kfold = *10*, the number of trees in the forest = *{500, 1000, 2000}*, learning rate = *{0.01, 0.1, 0.05},* the maximum depth of the tree = *{7, 9, 11, 15}*
MLP: with the method of GridSearchCV and kfold *= 10,* hidden layer size = *{{80, 80, 80}, {100, 100, 100}, {150, 150, 150}}, L2* penalty (regularization term) parameter = *{1e-3, 1e-4},* the solver for weight optimization = *{‘adam’, ‘sgd’},* tolerance for the optimization = *{1e-8, 1e-7, 1e-6}*
Gaussian NB: portion of the largest variance of all features that is added to variances for calculation stability = *{1e-16, 1e-14, 1e-12}*

### Evaluation

In order to evaluate the performance of the classifiers, the confusion matrices were used to compute sensitivity, specificity, accuracy and F-score (32, 87). Most researchers used weighted F-score to evaluate the classifier’s comprehensive performance. However, it leads to assessment bias between majority families and minority families. In this evaluation, we used macro F1 instead, which gives an arithmetic mean of the per-class F1-scores and avoids assessment bias to some extent. A statistical test was carried out in GraphPad Prism 8.3.0 using the Wilcoxon rank test and multiple Wilcoxon rank test at *p* < 0.05, 0.01, 0.001 level (“Wilcoxon rank test were performed using GraphPad Prism version 8.3.0 for Windows, GraphPad Software, La Jolla California USA, www.graphpad.com”).

We used the following abbreviations: True Positives (*TP*), False Positive (*FP*), True Negative (*TN)*, and False Negative (*FN*). The used formulas are as follows:

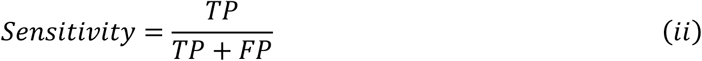

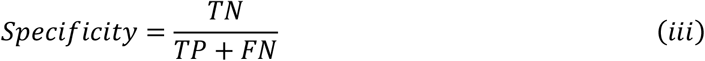

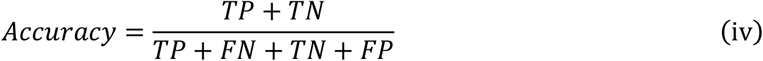

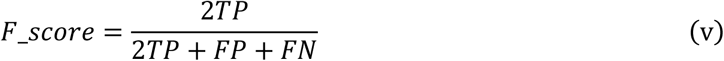

## ACKNOWLEDGMENTS

We would like to thank Ming Chen’s Bioinformatics Group members for assisting whenever needed. We also thank support given by the National Key Research and Development Program of China [2018YFC0310602; 2016YFA0501704]; National Natural Science Foundation of China [31571366, 31771477]; the Chinese Government Scholarship for foreign students(MOFCOM), Jiangsu Collaborative Innovation Center for Modern Crop Production, the Fundamental Research Funds for the Central Universities; and the Ministry of Education and Science of North Macedonia and the Ministry of Science and Technology (MOST) of China. Opinions, results, and conclusions articulated in this paper are those of the author(s) and do not necessarily reflect the views of the supporting organization.

## Author contributions

Solomon Shiferaw Beyene, Tianyi Ling, Blagoj Ristevski, and Ming Chen contributed equally and Solomon Shiferaw Beyene and Tianyi Ling write python code.

## Supplementary Table

**Supplementary Table S1.** The dataset used for the purpose of comparison of imbalanced and balanced datasets from Rfam database. The training (70%) and test dataset (30%) for classification and evaluation performance of machine learning algorithms. Feature distribution across different 16 riboswitch families using heat-map is shown in Figure 2.

| Type of Riboswitch family | Rfam ID | Full sequences | Training Set | Test set |
| --- | --- | --- | --- | --- |
| Purine Riboswitch | RF00167 | 813 | 587 | 226 |
| TPP riboswitch (THI element) | RF00059 | 3616 | 2551 | 1065 |
| SAM riboswitch (S box leader) | RF00162 | 1241 | 852 | 389 |
| Molybdenum Cofactor riboswitch | RF01055 | 303 | 210 | 93 |
| Glycine Riboswitch | RF00504 | 2125 | 1491 | 634 |
| Lysine Riboswitch | RF00168 | 341 | 245 | 96 |
| FMN Riboswitch | RF00050 | 1211 | 827 | 384 |
| Cobalamin riboswitch | RF00174 | 4286 | 2993 | 1293 |
| glmS glucosamine-6-phosphate activated ribozyme | RF00234 | 275 | 181 | 94 |
| SAM riboswitch (alpha-proteobacteria | RF00521 | 167 | 133 | 34 |
| yKoK leader | RF00380 | 217 | 152 | 65 |
| S-adenosyl methionine (SAM) riboswitch | RF00634 | 216 | 155 | 61 |
| PreQ1 Riboswitch | RF00522 | 116 | 81 | 35 |
| Cyclic di-GMP-I riboswitch(c-di-GMP-I) | RF01051 | 1020 | 701 | 319 |
| PreQ1-II (pre queuosine) riboswitch | RF01054 | 39 | 21 | 18 |
| S-adenosyl-L-homocysteine riboswitch | RF01057 | 198 | 148 | 50 |

**Supplementary Table S2.** Classifiers’ performances with balanced and imbalanced datasets arranged in F-score decreasing order in case of the balanced dataset. For a specific classifier, mean represents average sensitivity, specificity, accuracy and F-score value, while standard deviation (SD) depicted variation in different riboswitch families.

| Model | Accuracy |  | Specificity |  | Sensitivity |  | F-score |  |
| --- | --- | --- | --- | --- | --- | --- | --- | --- |
|  | Imbalanced | Balanced | Imbalanced | Balanced | Imbalanced | Balanced | Imbalanced | Balanced |
| **RF** | 0.996±0.004 | 0.997±0.001 | 0.925±0.118 | 0.972±0.011 | 0.997±0.003 | 0.998±0.001 | 0.945±0.083 | 0.975±0.049 |
| **MLP** | 0.997±0.004 | 0.997±0.001 | 0.958±0.068 | 0.969±0.012 | 0.998±0.004 | 0.998±0.001 | 0.961±0.057 | 0.970±0.042 |
| **SVM** | 0.996±0.005 | 0.996±0.001 | 0.943±0.059 | 0.964±0.014 | 0.997±0.005 | 0.997±0.001 | 0.955±0.049 | 0.968±0.050 |
| **GB** | 0.992±0.006 | 0.993±0.001 | 0.864±0.132 | 0.883±0.031 | 0.996±0.006 | 0.996±0.001 | 0.901±0.082 | 0.904±0.089 |
| **KNN** | 0.988±0.009 | 0.986±0.003 | 0.824±0.192 | 0.928±0.017 | 0.993±0.008 | 0.991±0.002 | 0.844±0.134 | 0.856±0.111 |
| **NB** | 0.979±0.013 | 0.985±0.002 | 0.814±0.117 | 0.841±0.028 | 0.989±0.014 | 0.991±0.002 | 0.705±0.212 | 0.771±0.163 |

**Supplementary Table S3.** The statistical difference of four measurements between the balanced and imbalanced datasets. Bolded *p*-values indicate the statistical difference (SD).

| Measurement | NB | MLP | RF | GB | SVM | KNN |
| --- | --- | --- | --- | --- | --- | --- |
| Accuracy | **0.0002** | **0.0096** | **0.0033** | **0.0326** | **0.0203** | 0.8169 |
| Sensitivity | **0.0496** | 0.118 | 0.1135 | 0.104 | **0.0173** | **0.0403** |
| Specificity | 0.1461 | **0.0038** | 0.3557 | 0.0977 | **0.0161** | 0.9305 |
| F-score | **0.0019** | 0.1406 | **0.0002** | 0.131 | **0.0008** | 0.95 |

